# Disruption of Myelin-Associated Glycoprotein Activity Drives Aberrant Cerebellar Neurodevelopment and Autism-Like Behaviors

**DOI:** 10.64898/2026.06.11.731430

**Authors:** Mara S. Mattalloni, Anabela Palandri, Glenda Martin Molinero, Cristian R. Bacaglio, Juan C. Molina, Alicia L. Degano, Pablo H.H. Lopez

## Abstract

Immune-mediated mechanisms have emerged as important contributors to neurodevelopmental vulnerability in a subset of autism spectrum disorder (ASD) cases. Circulating IgG antibodies against myelin-associated glycoprotein (MAG) have been reported in individuals with ASD and their mothers; however, the pathogenic relevance of these antibodies and the contribution of MAG signaling to neurodevelopment remain unclear. Here, we investigated whether disruption of MAG function during early postnatal life is sufficient to alter cerebellar development and induce ASD-relevant behavioral alterations. Using complementary genetic and immunological approaches, we show that constitutive deletion of *Mag* or transient postnatal blockade with a function-blocking anti-MAG antibody induces region-specific alterations in cerebellar development. MAG disruption results in excessive proliferation of granule cell precursors followed by delayed, region-restricted neuronal death, together with persistent abnormalities in Purkinje neuron number and dendritic maturation. These structural changes occur in the absence of major or persistent demyelination, consistent with dysregulation of myelin-associated developmental signaling rather than myelin loss. Importantly, early postnatal passive immunization with anti-MAG IgG is sufficient to induce significant impairments in social communication, sociability, and social recognition, recapitulating core behavioral domains relevant to ASD. Together, these data provide experimental evidence that immune-mediated interference with a myelin-associated signaling molecule can disrupt cerebellar development during critical postnatal windows and produce long-lasting behavioral consequences. Our findings identify MAG as a previously underappreciated regulator of neurodevelopment and support a model in which antibody-mediated perturbation of myelin-derived instructive signaling contributes to ASD-relevant phenotypes, aligning with emerging frameworks of immune-linked neurodevelopmental vulnerability.

**Highlights:**

- Disruption of myelin-associated glycoprotein (MAG) activity during early postnatal life alters cerebellar development in a region-specific manner.
- Genetic deletion or antibody-mediated blockade of MAG function induces granule cell dysregulation and Pkn structural abnormalities without overt demyelination.
- Passive immunization of anti-MAG antibodies is sufficient to induce autism-like behavioral phenotypes, supporting a causal immune-mediated mechanism.
- These findings identify myelin-associated signaling as a novel contributor to neurodevelopmental dysfunction and align with the maternal autoantibody–related ASD framework.

## Introduction

Autism spectrum disorders (ASD) comprise a heterogeneous group of neurodevelopmental conditions characterized by persistent impairments in social communication, restricted interests, repetitive behaviors, and altered sensory processing (Geschwind & Levitt, 2007; Lord et al., 2018; State & Šestan, 2012). Although genetic factors contribute substantially to ASD risk, heritability alone does not fully account for disease penetrance, phenotypic variability, or the marked sex bias observed in prevalence, supporting a multifactorial etiology involving non-genetic modifiers acting during critical windows of brain development (Courchesne et al., 2007; Geschwind, 2009; Sandin et al., 2017; State & Šestan, 2012).

Among these contributors, immune dysregulation and autoimmunity have emerged as relevant factors in a subset of individuals with ASD. Epidemiological studies report an increased prevalence of autoimmune diseases among first-degree relatives of individuals with ASD (Chen et al., 2016; Comi et al., 1999; Wills et al., 2007), while both affected children and their mothers harbor circulating IgG autoantibodies recognizing central nervous system (CNS) antigens, supporting a role for immune-mediated mechanisms in shaping neurodevelopmental trajectories (Diamond et al., 2009; Singer et al., 2006).

Experimental models of maternal immune activation further support this framework, as prenatal or early postnatal exposure to brain-reactive antibodies induces persistent neuroanatomical and behavioral alterations relevant to ASD (Estes & McAllister, 2015; Malkova et al., 2012; Patterson, 2011). However, the identity of the relevant antigenic targets and the mechanisms through which antibody-mediated perturbations disrupt neurodevelopment remain incompletely understood.

One particularly compelling, yet insufficiently explored, observation is the association between ASD and elevated titers of autoantibodies directed against myelin-associated glycoprotein (MAG). Several studies have reported increased levels of anti-MAG IgG antibodies in individuals with ASD, correlating with clinical severity, immune dysregulation, and increased prevalence of autoimmune disease in first-degree relatives (Mostafa et al., 2008; Mostafa & Al-Ayadhi, 2013; Vojdani et al., 2002). In addition, anti-MAG reactivity has been detected in both children with ASD and their mothers, supporting a potential contribution of maternal autoimmunity to neurodevelopmental risk (Abou-Donia et al., 2019). Despite these associations, whether anti-MAG antibodies exert a pathogenic role in ASD or represent an epiphenomenon remains unknown.

Beyond its traditional classification as a structural myelin protein, MAG is increasingly recognized as a biologically active signaling molecule at the axon–myelin interface. Although it represents only a minor component of central nervous system myelin, MAG exerts important regulatory functions in axon–glia communication, influencing neurite stability, neuronal survival, and postnatal circuit maturation (Lopez, 2014; Lopez & Báez, 2018). In this context, MAG enables myelin to exert instructive roles during postnatal brain maturation rather than functioning solely as a passive insulator. Consistently, MAG promotes motoneuron survival during early postnatal development and contributes to long-term axonal protection in both the central and peripheral nervous systems (Nguyen et al., 2009; Palandri et al., 2015).

The cerebellum represents a particularly relevant substrate for investigating the developmental consequences of MAG dysfunction. Once regarded primarily as a motor structure, the cerebellum is now recognized as a critical hub for cognitive, affective, and social processing, with strong anatomical and functional connections to cortical regions implicated in ASD (D’Mello & Stoodley, 2015; Fatemi et al., 2012; Wang et al., 2014). In particular, posterior cerebellar regions, including Crus I and Crus II, have been consistently associated with ASD-related abnormalities involving social cognition, behavioral regulation, and cerebro-cerebellar connectivity (Schmahmann, 1998; Stoodley et al., 2012; Wang et al., 2014; D’Mello et al., 2015; D’Mello & Stoodley, 2015; Wang et al., 2025).

In rodents, cerebellar development occurs predominantly during the early postnatal period and is characterized by tightly orchestrated waves of granule cell precursor proliferation, migration, differentiation, and apoptosis, alongside the maturation and refinement of Pkn dendritic architecture (Altman, 1997; Hatten & Heintz, 1995; Sotelo, 2004; Wang et al., 2014). This developmental window coincides precisely with the onset of active myelination and MAG expression, potentially rendering the cerebellum particularly vulnerable to perturbations in myelin-associated signaling pathways (Altman, 1997; D’Mello et al., 2015; D’Mello & Stoodley, 2015; Hatten & Heintz, 1995; Wang et al., 2014) Disruption of these coordinated developmental processes—particularly within cerebellar lobules linked to cognitive and affective functions—has been repeatedly associated with ASD-like phenotypes in both human studies and animal models (D’Mello et al., 2015; D’Mello & Stoodley, 2015; Fatemi et al., 2012; Wang et al., 2014).

Notably, the period of intense postnatal cerebellar remodeling overlaps with active myelination, suggesting that myelin-derived signals may contribute to neuronal maturation and circuit assembly (Fields, 2008; Fields et al., 2015; Mount & Monje, 2017). Within this developmental window, immune-mediated interference with MAG function could disrupt key neurodevelopmental processes without necessarily causing overt myelin defects. Consistently, combined deletion of MAG and Nogo-A impairs oligodendrocyte migration and delays myelination in several CNS regions, including the cerebellum (Pernet et al., 2008).

Collectively, these observations support the hypothesis that MAG dysfunction may link immune dysregulation to altered neurodevelopment relevant to ASD. Here, we tested whether genetic loss of MAG function or early postnatal antibody-mediated blockade is sufficient to induce ASD-relevant neuroanatomical and behavioral alterations. Using complementary genetic and immunological approaches, we examined the impact of MAG disruption on cerebellar development, neuronal homeostasis, and social behavior. Together, our findings provide experimental evidence linking immune-mediated disruption of MAG signaling to ASD-relevant neurodevelopmental phenotypes.

## Materials and Methods

### Animals

Mag-null founder mice were kindly provided by Dr. Bruce Trapp (The Cleveland Clinic Foundation, Cleveland, OH, USA). These mice were generated via targeted disruption of exon 5 of the *Mag* gene, as previously described (Li et al., 1994). To ensure genetic stability and >99% strain purity, mutant mice were repeatedly backcrossed onto a C57BL/6 background, with this process refreshed every two years (Pan et al., 2005).

Animals were randomly assigned to experimental groups based on age and genotype. Both male and female mice were utilized in the study; however, group assignment was conducted without explicit balancing for sex within each experimental cohort. Consequently, sex was treated as a random variable across age, genotype and treatment groups.

All procedures were conducted and sacrificed in accordance with the National Institutes of Health Guide for the Care and Use of Laboratory Animals and were approved by the Institutional Animal Care and Use Committee of the Instituto de Investigación Médica Mercedes y Martín Ferreyra (INIMEC-CONICET–Universidad Nacional de Córdoba) and later by the Institutional Animal Care and Use Committee of the Facultad de Ciencias Químicas, Universidad Nacional de Córdoba (RD-2026-299-E-UNC-DEC#FCQ).

### Passive immunization with anti-MAG monoclonal antibody

To disrupt MAG signaling during early postnatal development, wild-type pups received intraperitoneal injections of a function-blocking anti-MAG monoclonal antibody (IgG1, clone 513) (Meyer-Franke et al., 1995; Poltorak et al., 1987), kindly provided by Melitta Schachner (University of Heidelberg). C57BL/6 pups were administered daily injections of 50 µg anti-MAG antibody (1 mg/ml in 50 µl sterile saline) from postnatal day (P) 0 to P3, reaching a cumulative dose of 200 µg. Control littermates received equivalent injections of an isotype-matched monoclonal IgG (anti-c-Myc, clone 9E10; DSHB RRID:AB2266850). As with previous experiments, pups were randomly assigned to treatment groups regardless of sex, and experimental cohorts were not balanced for sex distribution. The hybridoma producing 9E10, originally developed by J. Michael Bishop, was obtained from the Developmental Studies Hybridoma Bank (University of Iowa, USA). This postnatal window was selected to coincide with early cerebellar development and the onset of myelination. Pilot experiments revealed no detectable differences between isotype IgG–treated and untreated wild-type mice; therefore, in accordance with reduction principles, untreated wild-type animals were used as reference controls in subsequent analyses.

### Reagents

The anti-MAG and anti-c-Myc mAbs were produced from hybridoma culture supernatants and purified using Protein A/G affinity chromatography, followed by dialysis against phosphate-buffered saline (PBS) as described (Lopez et al., 2010).

### Behavioral assays

#### Ultrasonic vocalizations

Ultrasonic vocalizations (USVs) were analyzed in offspring at postnatal day 7 (P7) and postnatal day 14 (P14). Animals were assigned to four experimental groups: (1) naïve control; (2) control IgG group; pups receiving intraperitoneal (i.p.) injections of control IgG on postnatal days P0–P3 (Wt group); (3) anti-MAG group, pups receiving i.p. injections of anti-MAG monoclonal antibody (1 mg/ml, 50 μl) on P0–P3; and (4) *Mag*-null mice. USVs were recorded using a paradigm adapted from previously described protocols (Crawley, 2007; Scattoni et al., 2008). Pups were removed directly from the home cage and temporarily separated from the dam while remaining with their littermates in the nest. Body temperature was monitored and maintained between 35–36.5 °C using a heating pad to avoid hypothermia-induced alterations in vocalization behavior.

Recordings were performed in a temperature-controlled, sound-attenuated chamber under dark conditions, as previously described (Yin et al., 2016). Individual pups were placed in the recording chamber and USVs were recorded for 5 min using an ultrasonic detector microphone (MedLab, Albany, USA) connected to Med-PC IV acquisition software. Vocalizations were analyzed in two ultrasonic frequency ranges (20–55 kHz and 55–100 kHz). After the initial recording session, pups were returned to the nest and subsequently isolated for 15 min before a second 5-min recording session. This procedure represents a modified maternal potentiation paradigm, in which the second separation typically elicits an increased number of calls compared with the first separation.

Maternal potentiation of USVs has been proposed to reflect early cognitive and communicative components of pup–mother interaction (Gal et al., 2023; Shair, 2007). Detected calls were visualized and quantified using Med-PC IV software, which automatically calculates acoustic parameters including total call number, call duration, and peak frequency.

#### Three-chamber social interaction test

The three-chamber social interaction test was performed at postnatal day 21 (P21), a developmental stage roughly comparable to early childhood in humans, when behavioral abnormalities associated with neurodevelopmental disorders become more apparent. Animals were assigned to the following experimental groups: naïve control (WT), pups injected ip. with control IgG on postnatal days P0–P3; anti-MAG group, pups injected ip. with anti-MAG antibody (1 mg/mL, 50 μL) on P0–P3; and *Mag*-null mice. The behavioral paradigm was adapted from the three-chamber sociability test originally described by Crawley et al. (2009) and widely used to evaluate sociability and social novelty preference in mice (Moy et al., 2004; Silverman et al., 2010). The apparatus consisted of a rectangular Plexiglas box divided into three compartments of equal dimensions (40.5 × 20 × 22 cm each). The side chambers communicated with the central chamber through openings (3.5 × 10 cm), allowing free movement between compartments. At the beginning of the session, the subject mouse was placed in the central chamber and allowed to habituate to the apparatus for 10 min with the doors closed. This was followed by an additional 10-min habituation period with the doors open, allowing free exploration of all three chambers. After habituation, the subject mouse was briefly confined to the central chamber while a novel unfamiliar mouse (stranger mouse) was placed in one of the side chambers under an inverted wire cup that allowed visual, olfactory, auditory, and limited tactile interaction. An empty inverted wire cup (novel object) was placed in the opposite chamber. Small weights were placed on top of the cups to prevent displacement by the subject mouse. The doors to the side chambers were then opened and the sociability test began, lasting 10 min. During this period, the following parameters were recorded: time spent in each chamber and number of entries into each chamber. Sociability was defined as the preference of the subject mouse to spend more time in the chamber containing the novel mouse than in the chamber containing the novel object. The number of chamber entries served as a control for general locomotor activity and exploratory behavior. To evaluate social novelty preference, an additional test was performed immediately afterward. A second unfamiliar mouse was placed in the chamber that was previously empty, and the time spent interacting with the newly introduced mouse versus the previously encountered mouse was recorded.

### Motor function assays

Body weight and motor performance were assessed to control for potential developmental or motor impairments that could confound the interpretation of social behavioral outcomes and to determine whether the experimental manipulations selectively affected social behavior in the absence of generalized motor deficits. Body weight and early developmental reflexes were evaluated at postnatal day 4 (P4). No significant differences in body weight were observed among experimental groups.

#### Negative geotaxis

Negative geotaxis was evaluated at postnatal day P4 as previously described (Crawley, 2004). Pups were placed head-down on a 45° inclined plane, and the latency required to rotate 180° and orient the head upward was recorded. Each animal was allowed two trials, and the latency to complete the maneuver was measured.

#### Postural (righting) reflex

The postural reflex was assessed as described (Crawley, 2007) by placing pups in a supine position on a flat surface and measuring the latency required to return to a prone position with all four paws contacting the surface. Each pup was allowed two attempts with a maximum duration of 15 seconds per trial. Righting within 3 seconds was considered indicative of reflex maturation.

### Rotarod performance

Motor coordination and balance were evaluated at postnatal day P21 using an accelerating rotarod apparatus (Rotamex 4/8 Rotarod, Columbus Instruments, OH, USA). Mice were placed individually on the rotating rod and subjected to an accelerating protocol in which the rod speed increased from 11 rpm to 27 rpm over 120 s, following parameters previously described for this apparatus. The rotarod consisted of a rotating rod (7.3 cm in diameter) positioned 28 cm above the base of the apparatus. Animals were carefully placed on the rod, and the latency to fall was recorded as a measure of motor coordination and balance. Each animal underwent four consecutive sessions on the rotarod. Each session lasted 4 minutes (240 s). When an animal fell from the rod, it was immediately returned to the apparatus to continue the session. The interval between sessions was 10 minutes, allowing sufficient recovery and minimizing fatigue between trials, consistent with previously described rotarod protocols in rodents. During each session, the following parameters were recorded: latency to first fall, total number of falls, and time between consecutive falls. From these measurements, the following variables were calculated: mean latency to first fall, mean number of falls per session, and average time between falls across sessions. These parameters were used as indices of motor coordination, balance, and motor learning. Animals were habituated to the apparatus prior to testing to reduce novelty-induced variability. Behavioral performance was analyzed using the mean values obtained across sessions for each parameter. Similar accelerating rotarod paradigms have been extensively used to assess motor coordination in mice (Rozas et al., 1997) and have been applied in behavioral characterization of mouse models of autism spectrum disorders (Peça et al., 2011).

### Histology and stereological analysis of cerebellar volume and granule cell number

Animals at P7, P14, and P21 were deeply anesthetized and transcardially perfused with PBS followed by 4% paraformaldehyde. Brains were post-fixed for 24 h at 4 °C, cryoprotected in 30% sucrose, embedded, and sectioned sagittally (30 µm) using a cryostat (Leica). Cresyl violet staining was performed for cytoarchitectural analysis of cerebellar architecture.

Cerebellar granule neuron (GC) populations were quantified at postnatal day 7 (P7) within the internal granular layer (IGL) of lobules VI and VII (primarily associated with cognitive processing) and lobule X (predominantly associated with motor function, serving as an internal control). Total cell numbers were estimated using unbiased design-based stereology via the Optical Fractionator method. Granule cells were identified based on their distinct nuclear morphology and their specific laminar position within the IGL.

Pkn populations were analyzed at postnatal day 14 (P14) in cerebellar lobules. Images were acquired using a light microscope, and individual lobules were reconstructed from multiple cerebellar sections obtained at 20× magnification (three serial sections per animal). Image reconstruction was performed using Adobe Photoshop CS6 (64-bit). Subsequent cell counting and morphometric analyses were conducted using Fiji software (ImageJ). Pkn were identified based on their characteristic large soma size and their localization within the Pkn layer.

All stereological analyses were performed using a computerized stereology workstation consisting of a modified light microscope (Olympus BX50 equipped with a PlanApo 1.25× objective [numerical aperture (N.A.) = 0.04] and a UPlanApo 20× oil-immersion objective [N.A. = 0.8]; Olympus, Tokyo, Japan), a motorized specimen stage for automated sampling along the X, Y, and Z axes (Ludl Electronics, Hawthorne, NY, USA), a CCD color video camera (HV-C20AMP; Hitachi, Tokyo, Japan), and stereology software (StereoInvestigator; MBF Bioscience, Williston, VT, USA). All stereological procedures were conducted by an investigator blinded to genotype and treatment groups to ensure unbiased data collection.

### Immunofluorescence

Free-floating sagittal brain cerebellar sections were blocked in PBS containing 5% normal donkey serum and 0.3% Triton X-100, followed by incubation with primary antibodies against Ki-67 (AB9260 Sigma-Aldrich; 1:500) and Calbindin-D-28K (Clone CB-955 Abcam; 1:200), according to the manufacturers’ instructions. Ki-67 immunostaining was used to label proliferating cells, as Ki-67 is expressed in all active phases of the cell cycle. Calbindin-D28K was used to label cell bodies and dendrite processes of Pkn. After washing, sections were incubated with appropriate Alexa-conjugated fluorescent secondary antibodies (Jackson ImmunoResearch Laboratories). When required, nuclei were counterstained with DAPI (4’,6-diamidino-2-phenylindole). Sections were mounted using Mowiol® 4-88 mounting medium. Fluorescence images were acquired with an Olympus Fluoview1000 confocal laser scanning microscope. To encompass the full thickness of the samples, serial optical sections (z-stacks) were captured at 5.0 µm intervals. The number of Ki-67+ and Calbindin+ cells was quantified within cerebellar lobules VI, VII, and X using Fiji (ImageJ) software. To ensure objectivity, all analyses were performed by an investigator blinded to the experimental groups.

### Fluoro-Jade C staining

Neurodegeneration was assessed using Fluoro-Jade C (FJC) (Millipore) staining as previously reported (Schmued et al., 2005). Briefly, 30 µm brain cerebellar sections were mounted on gelatin-coated slides (0.5% gelatin solution) and air-dried overnight. Sections were sequentially immersed in 100% ethanol (3 min), 70% ethanol (1 min), and distilled water (1 min), followed by incubation in 0.06% potassium permanganate solution for 10 min with gentle agitation. After rinsing in distilled water, sections were incubated in 0.0001% Fluoro-Jade C solution prepared in 0.1% acetic acid for 10 min in the dark.

Sections were rinsed in distilled water, air-dried, cleared in xylene, and coverslipped with DPX mounting medium. FJC-positive staining areas were visualized as bright green fluorescence under epifluorescence microscopy using appropriate FITC filter settings. Quantification of FJC-positive cells and stained area was performed in cerebellar lobules VI, VII, and X using Fiji (ImageJ) software. All analyses were conducted by an investigator blinded to experimental conditions.

### Statistical analysis

The statistical analyses were performed using GraphPad Prism 8.0.1. The statistical tests performed are described together with the results. In all cases, analysis of variance (ANOVA) was used and the number of ways depended on the number of factors and the design of each experiment. Values p <0.05 were considered statistically significant. In all cases, when a significant interaction was found (p < 0.05), a Tukey’ s test was performed as a post hoc analysis. All results were expressed as the mean ± standard error of the mean (SEM).

## RESULTS

### 1. MAG dysfunction induces region-specific alterations in early postnatal cerebellar architecture

To determine whether MAG expression influences postnatal cerebellar development, we performed a morphometric analysis of cerebellar structure in wild-type (WT), *Mag*-null mice and mice passively immunized with an IgG anti-MAG mAb (anti-MAG) at postnatal days P7, P14, and P21, encompassing the major stages of cerebellar maturation (**Figure 1**). Stereological analysis of cresyl violet–stained sections revealed a significant increase in volume from cerebellar lobules VI/VII, which are associated with cognitive and affective processing, in both *Mag*-null and anti-MAG–treated mice, whereas lobule X, predominantly linked to motor functions, displayed minor changes (**Figure 1A-B**). This difference was transient, as no significant volumetric changes were detected at P14 or P21.

**Figure 1.**
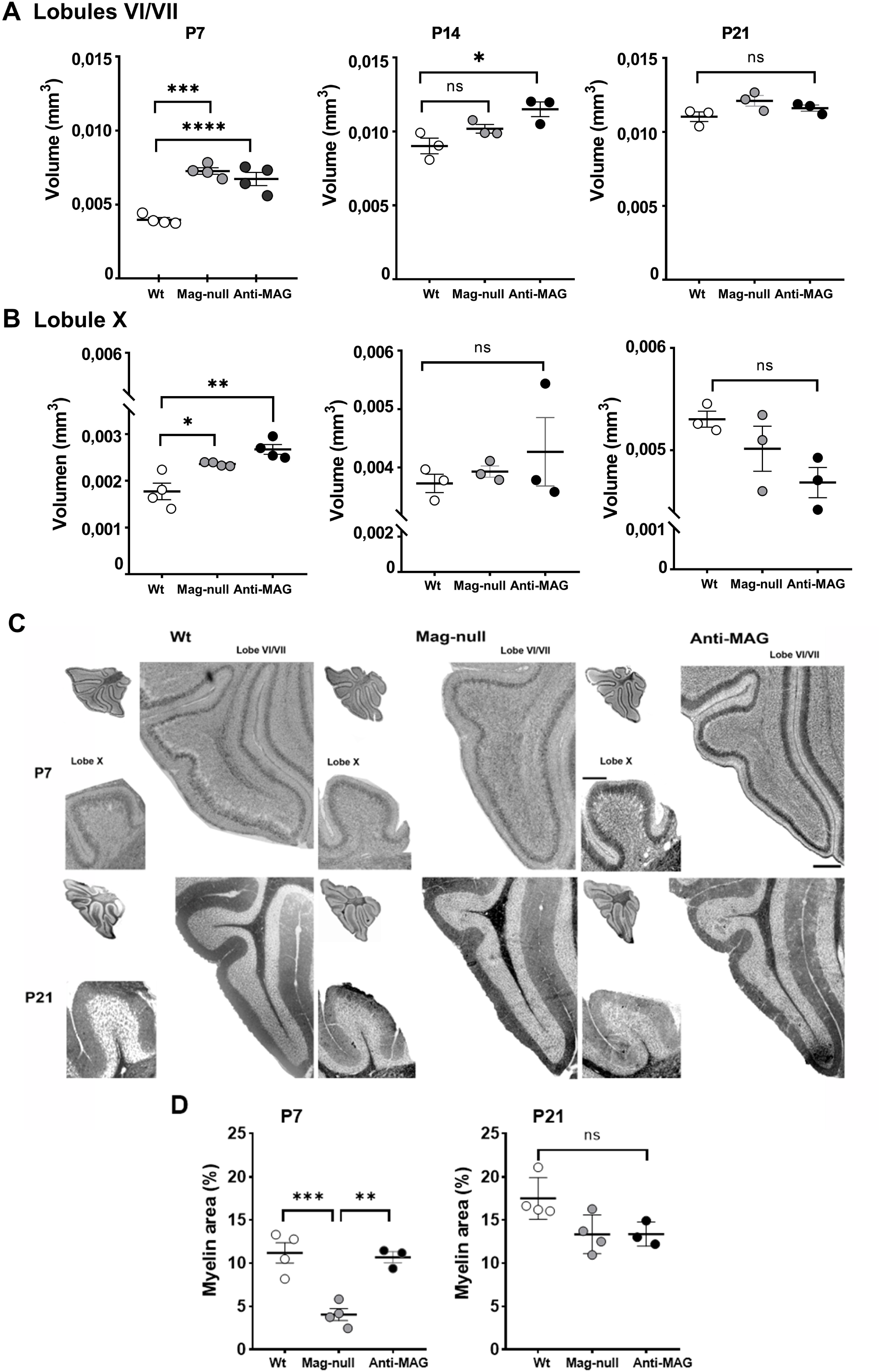
MAG disruption transiently alters cerebellar volume and myelin architecture. (A–B) Volumetric analysis of lobules **(A)** VI/VII and **(B)** X at postnatal days 7, 14, and 21. At P7, both *Mag-*null and anti-MAG mice exhibited enlarged lobules compared to WT. This increase persisted at P14 in Anti-MAG mice for lobules VI/VII, though all group differences resolved by P21. **(C)** Representative sagittal cerebellar sections stained with Sudan Black to visualize myelin; insets highlight lobules VI/VII and X. Scale bars: 400 µm (Lobes VI/VII); 200 µm (Lobes X). **(D)** Myelin area as a percentage of total tissue area: At P7, myelin content was significantly reduced in *Mag-*null mice versus WT; however, anti-MAG treatment did not significantly affect myelin area. By P21, these developmental differences were no longer detectable. Data are mean ± SEM; points represent biological replicates (n= 3-4 per group per time-point). Statistics: one-way ANOVA with Tukey’s post hoc test. \**p*< 0.05, \*\**p*< 0.01, \*\*\**p*< 0.001, \*\*\*\**p*< 0.0001; ns, not significant.

To determine if early cerebellar enlargement coincided with altered myelin development, we assessed myelination using Sudan Black staining at key postnatal stages. Qualitative analysis at P7 revealed a visible reduction in myelin labeling in both *Mag*-null and anti-MAG mice compared to WT controls, suggesting an initial delay in postnatal maturation (**Figure 1C**). However, quantitative analysis at P7 confirmed a significant decrease in Sudan Black–positive area specifically in *Mag*-null mice. In contrast, while anti-MAG mice exhibited a similar qualitative trend, their myelin density did not reach statistical significance for a deficit, despite these mice showing significant changes in overall cerebellar volume. By P21, myelin levels in both experimental groups increased progressively to reach levels comparable to WT controls (**Figure 1D**). These findings indicate that MAG deletion does not cause a permanent failure of myelination, but rather a transient maturation delay. Furthermore, MAG dysfunction selectively impacts early postnatal architecture in cerebellar regions linked to cognitive and social processing—sparing motor circuits—within a strictly defined developmental window.

### 2. MAG disturbance causes excessive granule cell precursor proliferation during early postnatal development

We next investigated whether changes in cerebellar volume were associated with alterations in granule cell (GC) populations, the most abundant neuronal population in mammalian brain. Unbiased stereological analysis of GC density in the internal granular layer (IGL) at P7 revealed a robust increase in GC numbers within lobules VI/VII in both *Mag*-null and anti-MAG, whereas lobule X showed no appreciable change. This regional difference persisted at P14 but resolved by P21, indicating a transient expansion of the GC population that coincided with changes in cerebellar volume (**Figure 2A-B**).

**Figure 2.**
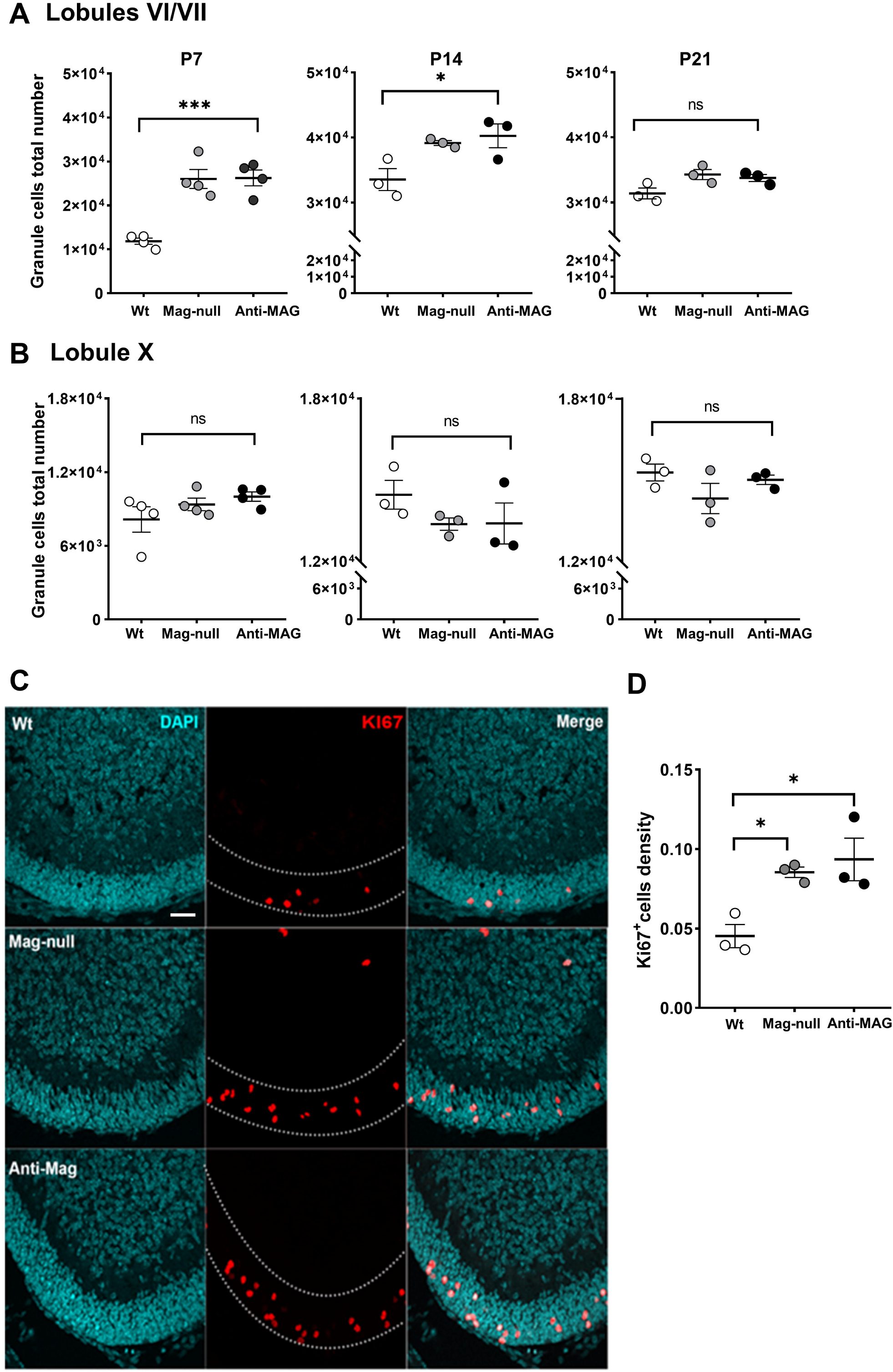
MAG disruption enhances GCP proliferation and transiently expands the granule cell population. (A–B) Estimated granule cell (GC) populations in lobules **(A)** VI/VII and **(B)** X at P7, P14, and P21. Both *Mag-*null and anti-MAG mice exhibited increased GC numbers at early stages, with populations normalizing to WT levels by P21. GC populations were estimated via stereological analysis (Stereo Investigator). **(C)** Representative confocal images of cerebellar sections at P7 immunostained for Ki-67 (red) and counterstained with DAPI (cyan). Dotted lines delineate the external granule layer (EGL); merged images highlight proliferating granule cell precursors (GCPs). Scale bar: 30 µm. **(D)** Proliferation in the EGL from lobules VI/VII at P7; quantification of Ki-67^+^ cell density across experimental groups. Values were normalized to the EGL area and expressed as cells/100 µm^2^. Both *Mag-*null and anti-MAG animals exhibited significantly increased proliferative activity compared to WT. Data are expressed as mean ± SEM; points represent biological replicates (n= 3-4 per group per time-point). Statistics: one-way ANOVA with Tukey’s post hoc test. *\*p <* 0.05*, ***p<* 0.001; ns, not significant.

Given the increased GC density associated with MAG deficiency, we next examined whether this phenotype stems from altered granule cell precursor (GCP) proliferation. Immunofluorescence staining for the marker Ki-67 at P7—the peak of mitotic activity—revealed a significant increase in proliferating cells within the External Granule Layer (EGL) of both *Mag*-null and anti-MAG groups relative to WT controls **(Figure 2C**). Interestingly, quantitative analysis showed that hyper proliferation was topographically restricted to lobules VI/VII; conversely, proliferation rates in lobule X remained comparable across all groups **(Figure S1)**. Together, these data suggest that MAG-dependent signaling is a key regulator of GCP proliferation during a critical window of cerebellar development.

### 3. MAG Activity is required for the timely elimination of excess granule cells

The transient nature of increased GC density suggested that excess granule neurons generated early in development may subsequently undergo cell death. To test this hypothesis, we assessed neurodegeneration in cerebellum using Fluoro-Jade C (FJC) staining at P7, P14, and P21 (**Figure 3A**). Both *Mag*-null and anti-MAG mice exhibited a robust increase in FJC-positive labeling within the IGL at P14 and P21—overlapping with the period of physiological apoptosis of redundant GC. Importantly, this elevation was restricted to lobules VI/VII, whereas lobule X showed increased staining only in P14 *Mag*-null mice (**Figure 3C**). No differences in FJC staining were observed at P7 among the groups, suggesting that neurodegeneration occurs after the initial proliferative phase (**Figure 3B, C**). Together, these results suggest that MAG disruption uncouples the normal balance between GC production and developmental-driven programmed cell death.

**Figure 3.**
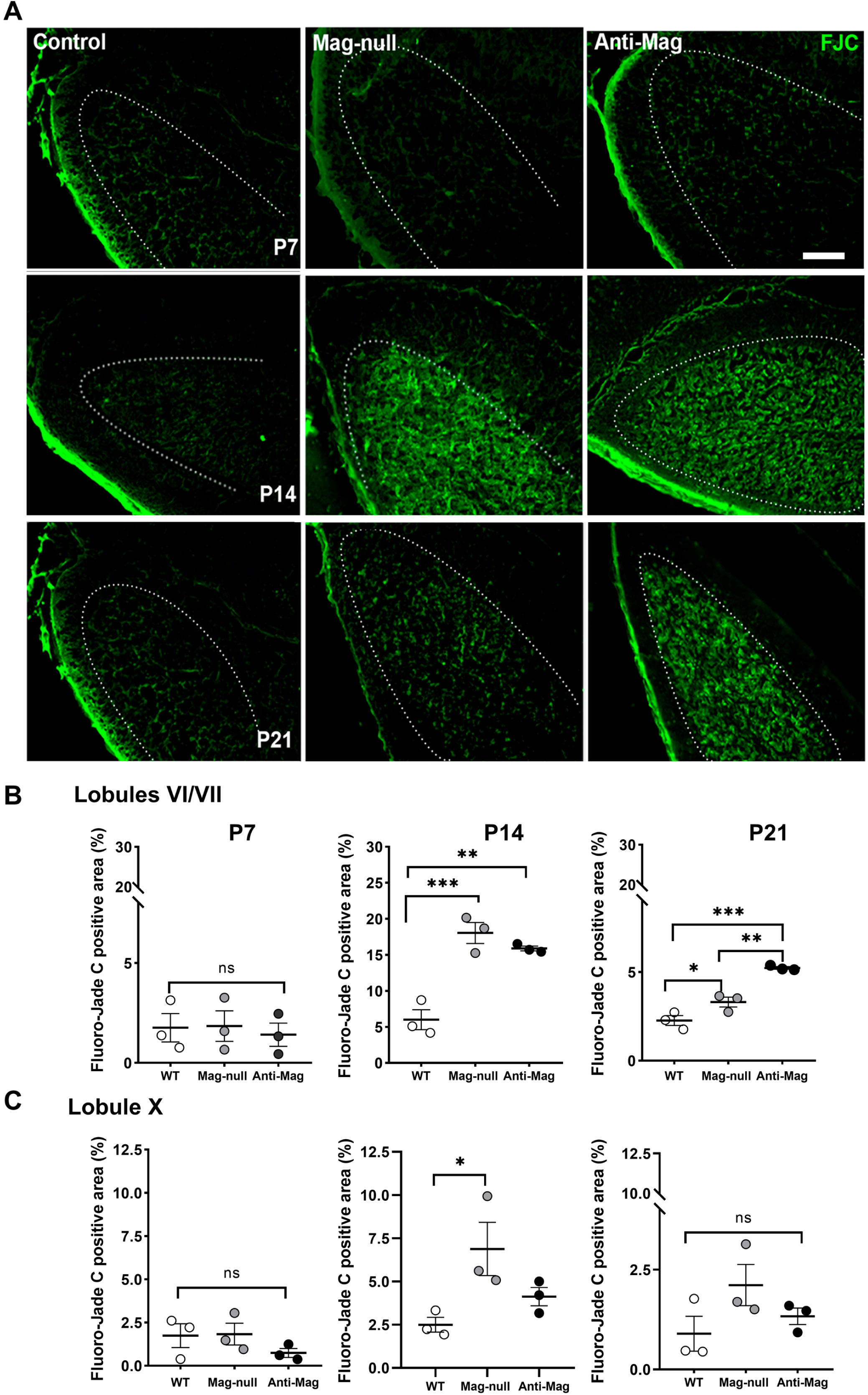
MAG disruption triggers delayed neurodegeneration of excess granule cells. **(A)** Representative sagittal sections of the cerebellum stained with Fluoro-Jade C (FJC) at P7, P14, and P21 to identify degenerating neurons. Scale bar: 100 µm. **(B–C)** Quantification of FJC-positive area in lobules **(B)** VI/VII and **(C)** X at P7, P14, and P21. Both *Mag*-null and anti-MAG-treated mice exhibited a robust increase in FJC labeling within the internal granule layer (IGL) of lobules VI/VII at P14 and P21. In lobule X, significantly elevated FJC staining was restricted to *Mag*-null mice at P14; no significant differences were observed across groups at P7. Data are mean ± SEM; points represent biological replicates (n= 3-4 per group per time-point). Statistics: one-way ANOVA with Tukey’s post hoc test. \**p* < 0.05, \*\**p* < 0.01, \*\*\**p*< 0.001, ns, not significant.

### 4. MAG disruption alters Purkinje cell number and dendritic development in cognitive cerebellar lobules

Given the central role of Purkinje cells (Pkn) in cerebellar circuit assembly and ASD-related phenotypes (Fatemi et al., 2002; Skefos et al., 2014), we examined whether MAG disruption/deletion impacts Pkn development. Immunofluorescence analysis using anti-calbindin antibodies were performed at P7 and P14 (**Figure 4 A, B**). Both *Mag*-null and anti-MAG groups exhibited a significant increase in total Pkn numbers compared to WT controls (**Figure 4 C, D**), a phenotype particularly pronounced in lobules VI/VII while lobule X remained unaffected (**Figure S2**). Beyond quantitative changes, MAG-deficient Pkn displayed distinct morphological abnormalities; at P7, somata appeared elongated and atypical, while dendritic arbors were significantly shorter and less complex than those in WT mice (**Figure 4A, E; Figure S2**). These impairments in both cell density and dendritic architecture persisted through P14, indicating that MAG signaling is essential for the early positioning, survival, and structural maturation of Purkinje cells.

**Figure 4.**
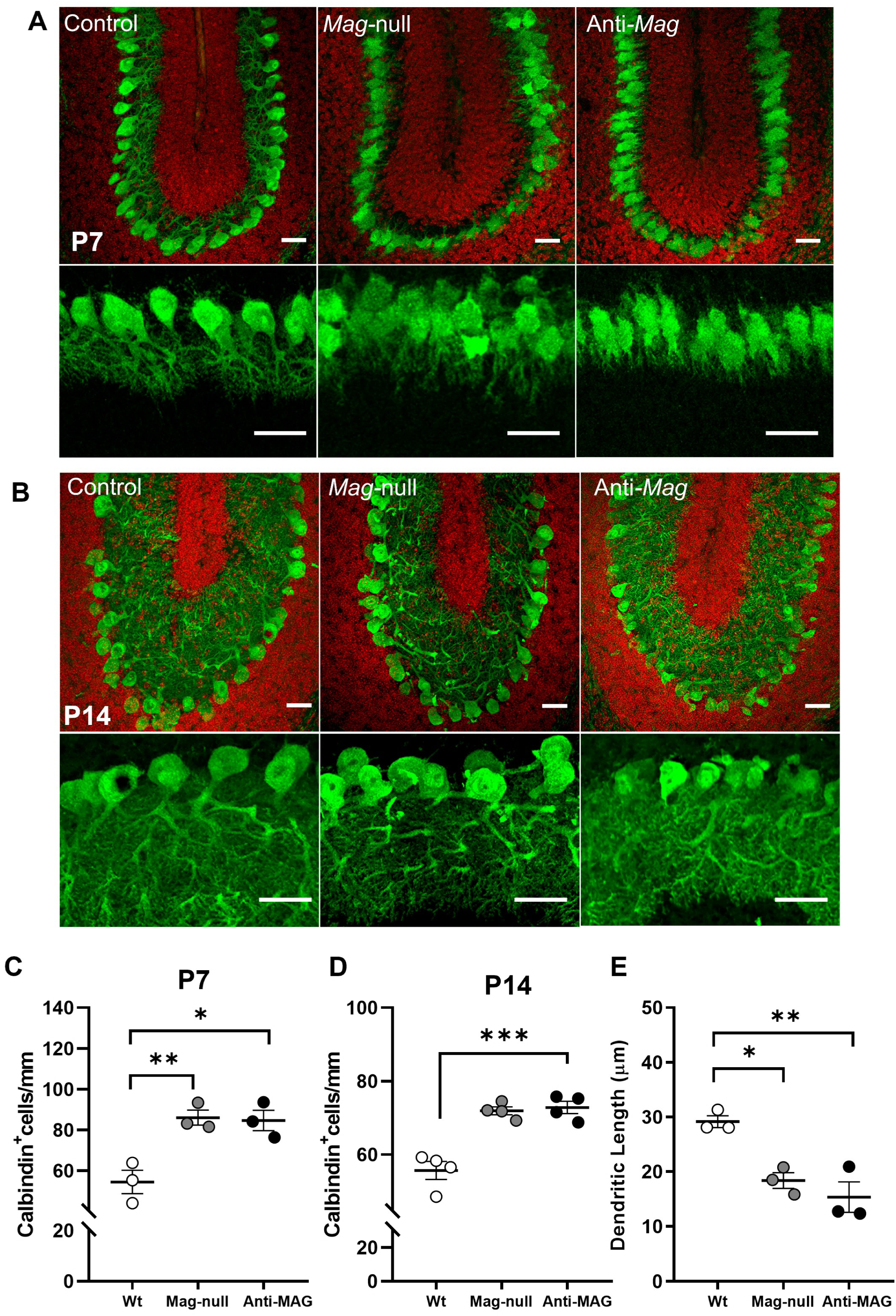
MAG disruption increases Purkinje cell (Pkn) population and impairs dendritic development. (A, **B)** Representative confocal images of cerebellar lobule VI/VII sections from WT, *Mag*-null, and anti-MAG groups immunostained for Calbindin (green) and DAPI (red) at P7 and P14, respectively. Images illustrate Pkn morphology, dendritic arborization within the molecular layer, and alignment along the Pkn layer. **(C, D)** Quantitative analysis reveals a persistent increase in Calbindin-positive Pkn numbers in both *Mag*-null and anti-MAG groups compared to WT. **(E)** Measurement of dendritic length at P7 shows significantly shorter arbors in MAG-deficient groups relative to WT. Data represent mean ± SEM (n = 4 per group). Statistical significance was determined by one-way ANOVA with Tukey’s post hoc test; \*\**p* < 0.01, \*\*\*\**p*< 0.0001, ns, not significant. Scale bars: 30µm.

### 5. Early disruption of MAG signaling impairs social communication without affecting motor reflexes

To determine whether early cerebellar anatomical alterations induced by MAG disruption are correlated with altered behavior, we first assessed social communication in neonatal mice using ultrasonic vocalization (USV) analysis at P7. Both *Mag*-null and anti-MAG groups exhibited a significant reduction in the number of USVs compared with their respective control groups (**Figure 5A**). This reduction was observed across both low-frequency (20–55 kHz) and high-frequency (55–100 kHz) call categories, which are associated with aversive/defensive and affiliative communication, respectively. Notably, maternal potentiation testing led to a convergence of phenotypes across experimental groups with the exception of *MAG*-null mice which displayed increased USVs at 20-55 kHz compared to control, indicating preserved vocalization capacity and arguing against motor or respiratory deficits (**Figure 5B**).

**Figure 5.**
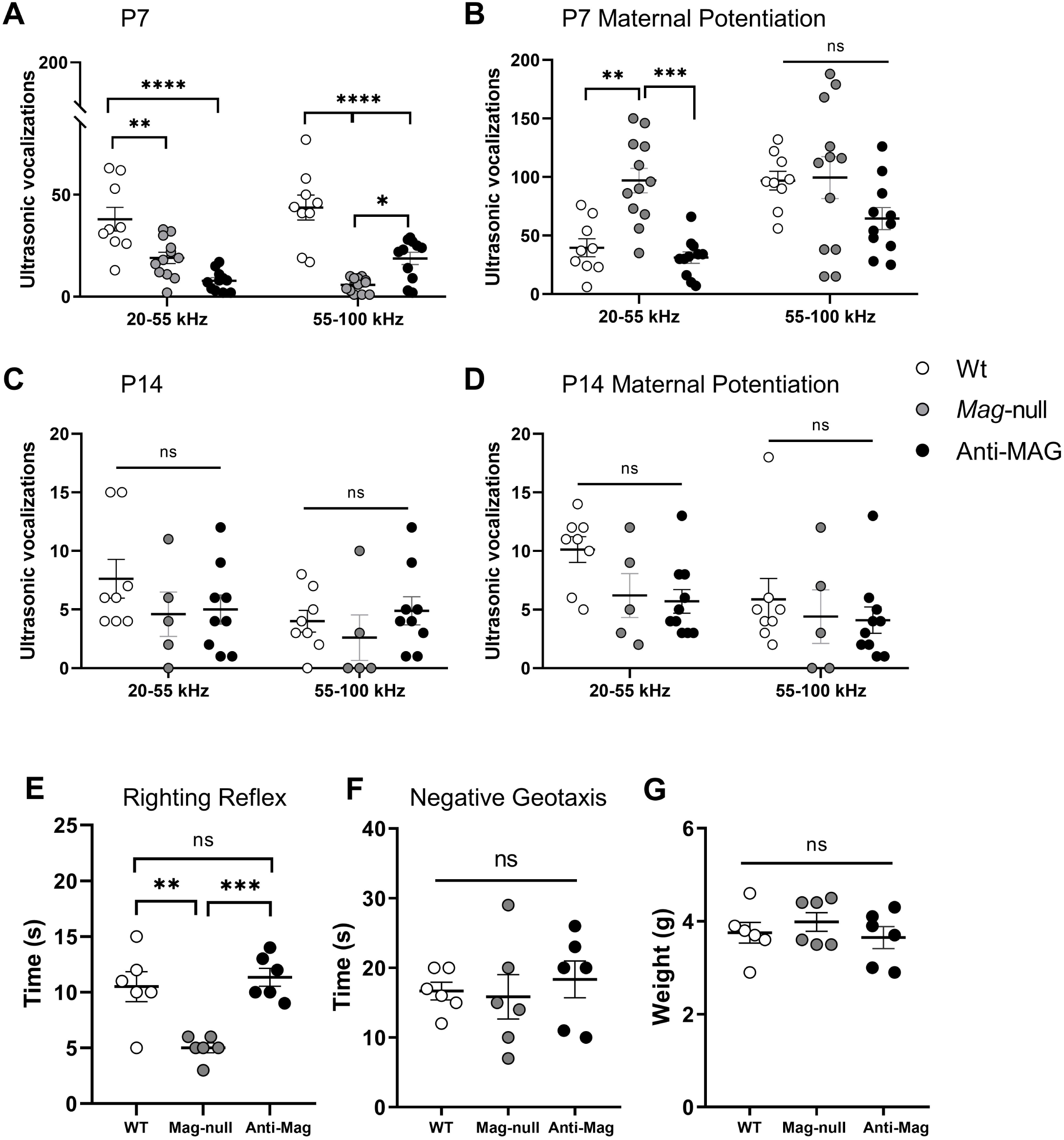
MAG disruption alters early developmental reflexes and ultrasonic vocalization behavior. These panels illustrate early developmental assays evaluating motor and communicative behaviors across experimental groups during the postnatal period. Quantification of ultrasonic vocalizations (USVs) at P7 without **(A)** or with maternal potentiation **(B)** revealed significant group-dependent differences in call frequency and acoustic profiles. **(C, D)** Conversely, no significant differences in vocalization patterns were observed at P14. Together, these data indicate that MAG deficiency selectively alters early communicative behavior during neonatal development. **(E–F)** Assessment of negative geotaxis and righting reflex performance demonstrated subtle delays in motor maturation only in Mag-null animals compared with WT. **(G)** No significant changes in body weight were observed among the groups. Data are presented as mean ± SEM with individual data points representing biological replicates (n=6-12 per group). Statistical comparisons were performed using two-way ANOVA (**A-D**) or one-way ANOVA (**E-G**) followed by Tukey’s post hoc test. Significance is indicated as \**p*< 0.05, \*\**p*< 0.01, \*\*\**p*< 0.001, \*\*\*\**p*< 0.0001; ns, not significant.

In contrast, no significant differences in USVs emission were observed at P14 among experimental groups (**Figure 5C, D**). Rather than indicating a transient effect, this likely reflects the distinct developmental roles of USVs, which peak during early neonatal stages (e.g., P7), prior to eye opening, and decline markedly thereafter as sensory modalities mature and communicative demands change.

Assessment of early motor reflexes, including negative geotaxis and postural reflexes, revealed no significant differences between anti-MAG-treated and control mice, indicating that antibody-mediated MAG disruption selectively impairs social communication while sparing basic motor function. In contrast, Mag-null mice showed normal performance in the negative geotaxis test but exhibited enhanced postural reflex responses compared to control animals. (**Figure 5E, F**). In addition, no differences in body weight were detected between groups at P7 (**Figure 5G**), further supporting the absence of general motor developmental delay.

### 6. MAG inactivation induces autism-like sociability and social recognition deficits at P21

We next examined whether the early disruption of MAG signaling leads to persistent behavioral alterations during later developmental stages. Sociability and social memory were assessed at P21 using the three-chamber social interaction test (**Figure 6 A-C**). This developmental time point represents a stage where major cerebellar architectural features are finalized and GC migration has largely subsided (Butts et al., 2014; Sillitoe & Joyner, 2007; White & Sillitoe, 2013). Both anti-MAG and *Mag*-null mice exhibited markedly diminished sociability, characterized by a significant reduction in the time spent exploring a novel conspecific compared to WT controls (**Figure 6D**).

**Figure 6.**
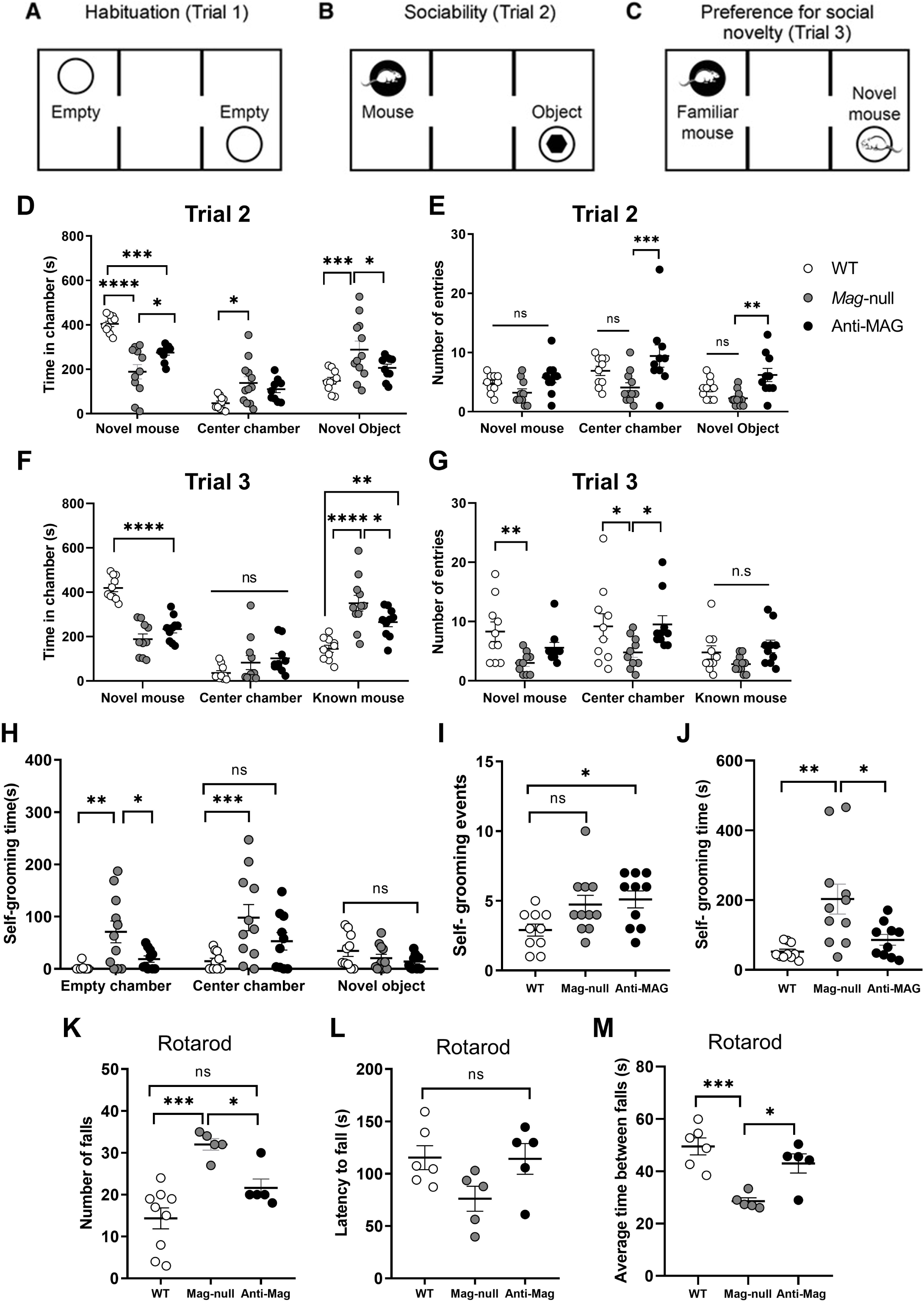
MAG disruption alters social behavior and exploratory responses. (A–C) Schematic representation of the behavioral paradigms used to assess social interaction, including habituation **(A)**, sociability **(B)**, and social novelty preference **(C)**. These tasks were performed in WT, *Mag*-null and anti-MAG mice. **(D–E)** Behavioral performance during the sociability phase (Trial 2) revealed group-dependent differences in exploratory activity and chamber preference, indicating altered social approach behavior in MAG-deficient animals. **(F–G)** Analysis of social novelty preference (Trial 3) demonstrated differences in the exploration of familiar versus novel conspecifics across groups. **(H–M)** Additional behavioral measures, including repetitive grooming and motor coordination (rotarod), further supported altered behavioral performance associated with MAG disruption. Data are presented as mean ± SEM with individual data points representing biological replicates (n=6-12 per group). Statistical comparisons were performed using one-way ANOVA followed by Tukey’s post hoc test. Significance levels were defined as \**p* < 0.05, *\*\*p* < 0.01, *\*\*\*p* < 0.001, *\*\*\*\*p <* 0.0001; ns, not significant.

During the social recognition test, WT controls demonstrated a significant preference for the novel stimulus, as expected. In contrast, both anti-MAG and *Mag*-null mice exhibited a marked avoidance of the novel stimulus, showing a preference for the familiar subject (**Figure 6F**). This atypical social behavior indicates an impairment in standard social recognition memory and suggests a potential shift toward social anxiety or neophobia. Notably, these behavioral deficits occurred without significant differences in the total number of chamber entries in both MAG deficient groups, indicating that the observed social impairments are not confounded by altered locomotor activity or a lack of exploratory drive (**Figure 6 E, G**).

Self-grooming behavior was analyzed in parallel with social testing within the same arena. Quantitative analysis revealed distinct grooming phenotypes between the groups: *Mag*-null mice exhibited an increase in total grooming duration (**Figure 6J**), particularly within the empty and central chambers (**Figure 6H**), despite a stable frequency of grooming bouts (**Figure 6I**). In contrast, the anti-MAG group showed a significant elevation in bout frequency without a corresponding change in total grooming time, an effect that persisted regardless of chamber location (**Figure 6H–J**). These behavioral abnormalities resemble core features of ASD-like phenotypes described in several rodent models of neurodevelopmental disorders.

### 7. Motor coordination is differentially affected in MAG-null and anti-MAG mice

Finally, motor coordination was assessed using the rotarod test at P21. Anti-MAG mice performed comparably to control mice across all measured parameters, including latency to first fall and total number of falls. In contrast, *Mag*-null mice exhibited a modest but significant impairment in motor coordination, characterized by an increased number of falls and reduced performance over repeated trials (**Figure 6K-M**), in accordance with a previous report (Pan et al., 2005).

These findings indicate that transient postnatal blockade of MAG signaling is sufficient to induce persistent social and cognitive behavioral deficits in the absence of major motor impairment, whereas constitutive MAG deletion produces broader motor consequences.

## DISCUSSION

In this study, we provide experimental evidence that disruption of MAG activity during early postnatal life is sufficient to induce region-specific neurodevelopmental alterations in the cerebellum and persistent ASD-relevant behavioral phenotypes. Using complementary genetic and immunological approaches, we demonstrate that both constitutive MAG deletion and transient postnatal blockade of MAG function converge on a common developmental trajectory characterized by abnormal cerebellar morphogenesis, dysregulated GC homeostasis, and alterations in Pkn number and dendritic maturation, together with selective impairments in social communication and sociability. These findings support a causal contribution of MAG dysfunction to neurodevelopmental outcomes relevant to ASD and provide proof of concept for an immune-linked neurodevelopmental pathway affecting cerebellar circuit assembly (Becker & Stoodley, 2013; Fatemi et al., 2012; S. S.-H. Wang et al., 2014).

Our data identify MAG as a critical modulator of cerebellar development during a narrow postnatal window. Rather than reflecting static changes in cell number, the observed phenotype follows a dynamic sequence, including transient expansion of cerebellar volume at P7, excessive proliferation of GCP in lobules VI and VII, and delayed, region-restricted elimination of granule neurons at later stages. This pattern suggests disruption of homeostatic mechanisms coordinating neuronal production, maturation, and elimination (Altman, 1997; Hatten & Heintz, 1995; Wang & Zoghbi, 2001).

Development of Pkn was also differentially affected. Although largely spared from cell death, they exhibited increased numbers and persistent alterations in dendritic morphology, consistent with impaired structural maturation. Given that Pkn represent the sole output neurons of the cerebellar cortex, even subtle perturbations in their differentiation are likely to have disproportionate consequences for circuit integration and function (Palay, 1974; Tsai et al., 2012). Notably, Pkn abnormalities are among the most consistent neuropathological findings in ASD (Fatemi et al., 2002; Skefos et al., 2014), supporting the translational relevance of these observations.

One possible mechanistic framework involves signaling pathways downstream of MAG, including RhoA-associated pathways previously implicated in neuronal survival and cerebellar development. MAG activates the p75/NgR/GT1b/Lingo-1 receptor complex, leading to engagement of the RhoA/ROCK pathway and regulation of cytoskeletal dynamics, neurite stability, and neuronal survival (Domeniconi et al., 2002; Lopez & Báez, 2018; Mi et al., 2004; Yiu et al., 2006). Importantly, the consequences of RhoA activation are highly context-dependent: in early postnatal motoneurons, this pathway promotes survival (Palandri et al., 2015), whereas in GC it can trigger apoptosis (Fernández-Suárez et al., 2019; Semenova et al., 2007). Consistently, genetic and pharmacological studies show that both excessive and insufficient RhoA/ROCK signaling disrupt cerebellar development, leading to defects in lamination, migration, and neuronal survival (Govek et al., 2005; Mulherkar et al., 2014). Together, these observations suggest that disruption of MAG-dependent signaling may alter developmental thresholds controlling neuronal survival and elimination. In this context, the structural and behavioral phenotypes observed here can be interpreted as the consequence of subtle shifts in signaling balance rather than overt neurodegeneration.

Importantly, our findings should be distinguished from classical anti-MAG–associated demyelinating disorders. In contrast to adult-onset anti-MAG neuropathies characterized by progressive demyelination (Dalakas, 2010), our model does not induce overt myelin loss. Instead, the phenotype is consistent with dysregulation of myelin-associated signaling during a critical developmental window. Although MAG represents a minor quantitative component of myelin, it plays a major signaling role at the axon–glia interface (Lopez, 2014). Disruption of MAG function during early postnatal life is therefore unlikely to destabilize mature myelin, but may alter the timing and quality of myelin-dependent signals required for neuronal maturation and circuit refinement (Almeida & Lyons, 2017; Fields, 2008).

This interpretation is further supported by the developmental timing of myelination, which overlaps extensively with processes such as neuronal migration, differentiation, apoptosis, and synaptogenesis (Almeida & Lyons, 2017; Brody et al., 1987; Fields, 2008; Yakovlev, 1967). Within this framework, even transient perturbations in myelin-associated signaling may be sufficient to redirect developmental trajectories without producing classical demyelinating pathology. Consistently, altered MAG expression has been reported in mouse models of ASD-related genes such as SLC25A12 (Sakurai et al., 2010), suggesting that myelin-associated signaling pathways may be selectively vulnerable to neurodevelopmental perturbations.

An important strength of the present study lies in the use of passive antibody transfer to interrogate the pathogenic potential of anti-MAG IgG. This strategy parallels maternal autoantibody–related ASD (MAR-ASD) models, in which antibody exposure alone is sufficient to induce long-lasting neuroanatomical and behavioral abnormalities in the absence of inflammation (Braunschweig et al., 2012; Brimberg et al., 2013). Although our study does not directly address prenatal exposure, the effects observed following early postnatal MAG blockade are conceptually aligned with these models, as both involve antibody-mediated interference with developmentally relevant signaling pathways rather than structural tissue damage (Diamond et al., 2009).

Notably, both MAR-ASD models and our experimental paradigm converge on the cerebellum—particularly lobules implicated in cognitive and social processing—as a preferential target of antibody-mediated developmental disruption (Becker & Stoodley, 2013; Martínez-Cerdeño et al., 2016). In addition, antibodies directed against MAG have been identified in children with ASD (Mostafa et al., 2008; Mostafa & Al-Ayadhi, 2013) and, in both ASD children and their mothers (Brimberg et al., 2013), supporting the possibility that MAG-directed immunity may contribute to neurodevelopmental vulnerability. While the temporal origin and pathogenic relevance of these antibodies remain to be fully established, our findings provide experimental proof of concept that antibody-mediated disruption of MAG signaling is sufficient to alter cerebellar maturation and produce long-lasting behavioral consequences.

In a more speculative context, anti-MAG autoimmunity may arise through mechanisms such as molecular mimicry, particularly given that these antibodies recognize carbohydrate epitopes including sulfated glucuronyl paragloboside (SGPG) (Latov, 2014; Steck, 2021). Although this hypothesis remains to be tested in ASD, it provides a biologically plausible framework for future investigation.

Several limitations should be acknowledged. First, the downstream molecular mechanisms linking MAG disruption to the observed structural changes remain to be defined. Second, behavioral analyses focused on core ASD-relevant domains and may not capture the full spectrum of cerebellar-related phenotypes. Third, our investigation was restricted to the cerebellum; this anatomical focus was guided by well-documented cerebellar alterations in ASD patients (Bauman & Kemper, 2005; Fatemi et al., 2012), and the region’s established role in the disorder’s behavioral manifestations (D’Mello & Stoodley, 2015; Wang et al., 2014), though it leaves potential extra-cerebellar contributions unexplored. Finally, in order to minimize animal usage while establishing the primary phenotype, sex-specific effects were not addressed in this round of studies, despite known differences in ASD prevalence and cerebellar development.

In summary, our study identifies MAG as a critical regulator of postnatal cerebellar development and demonstrates that immune-mediated disruption of MAG signaling is sufficient to induce persistent ASD-related phenotypes. These findings support a model in which myelin-associated signaling pathways act as instructive regulators of neurodevelopment and highlight antibody-mediated modulation of axon–glia signaling as a potential mechanism contributing to ASD heterogeneity. Future studies will be required to determine the clinical relevance of anti-MAG autoimmunity and to assess whether targeting myelin-associated signaling pathways may offer therapeutic opportunities in defined patient subsets.

**Figure S1. GCP proliferation is not altered after MAG disruption in cerebellar lobule X**. **(A)** Representative confocal images of lobule X cerebellar sections at P7 immunostained for Ki-67 (red) and counterstained with DAPI (cyan). Dotted lines delineate the external granule layer (EGL); merged images highlight proliferating granule cell precursors (GCPs). Scale bars: 30µm. **(B)** Quantification of Ki-67^+^ cell density across experimental groups. Values were normalized to the EGL area and expressed as cells/100 µm^2^. There were no observable differences in proliferative activity between WT, *Mag*-null, and anti-MAG animals. Data are expressed as mean ± SEM; points represent biological replicates (n= 3-4 per group per time-point). Statistics: one-way ANOVA with Tukey’s post hoc test. ns, not significant.

**Figure S2. MAG disruption impairs Purkinje (Pkn) cell morphology and arborization in lobule X without reducing cell numbers. (A, B)** Representative confocal images of lobule X cerebellar sections from WT, *Mag*-null, and anti-MAG groups immunostained for Calbindin (green) and DAPI (red) at P7 (A) and P14 (B). Images depict abnormal Pkn morphology and shorter dendritic arborizations within the molecular layer at P7 in MAG-deficient groups relative to WT. Scale bars: 30µm. **(C, D)** Quantitative analysis revealed no significant differences in the number of Calbindin-positive Pkn cells among the WT, *Mag*-null, and anti-MAG groups at either P7 or P14. Data represent mean ± SEM (n = 4 per group). Statistical significance was determined by one-way ANOVA with Tukey’s post hoc test; ns, not significant.

## References

1. Abou-Donia, M. B., Suliman, H. B., Siniscalco, D., Antonucci, N., ElKafrawy, P., & Brahmajothi, M. V. (2019). de novo Blood Biomarkers in Autism: Autoantibodies against Neuronal and Glial Proteins. Behavioral Sciences, 9(5), 47. 10.3390/bs9050047.

2. Almeida, R. G., & Lyons, D. A. (2017). On Myelinated Axon Plasticity and Neuronal Circuit Formation and Function. Journal of Neuroscience, 37(42), 10023–10034. 10.1523/JNEUROSCI.3185-16.2017.

3. Altman, J. (with Bayer, S. A.). (1997). Development of the cerebellar system: In relation to its evolution, structure, and functions. CRC Press.

4. Bauman, M. L., & Kemper, T. L. (2005). Neuroanatomic observations of the brain in autism: A review and future directions. International Journal of Developmental Neuroscience, 23(2-3), 183–187. 10.1016/j.ijdevneu.2004.09.006.

5. Becker, E. B. E., & Stoodley, C. J. (2013). Autism spectrum disorder and the cerebellum. International Review of Neurobiology, 113, 1–34. 10.1016/B978-0-12-418700-9.00001-0.

6. Braunschweig, D., Golub, M. S., Koenig, C. M., Qi, L., Pessah, I. N., Van De Water, J., & Berman, R. F. (2012). Maternal autism-associated IgG antibodies delay development and produce anxiety in a mouse gestational transfer model. Journal of Neuroimmunology, 252(1-2), 56–65. 10.1016/j.jneuroim.2012.08.002.

7. Brimberg, L., Sadiq, A., Gregersen, P. K., & Diamond, B. (2013). Brain-reactive IgG correlates with autoimmunity in mothers of a child with an autism spectrum disorder. Molecular Psychiatry, 18(11), 1171–1177. (91616345). 10.1038/mp.2013.101.

8. Brody, B. A., Kinney, H. C., Kloman, A. S., & Gilles, F. H. (1987). Sequence of Central Nervous System Myelination in Human Infancy. I. An Autopsy Study of Myelination. Journal of Neuropathology & Experimental Neurology, 46(3), 283–301. 10.1097/00005072-198705000-00005.

9. Butts, T., Green, M. J., & Wingate, R. J. T. (2014). Development of the cerebellum: Simple steps to make a ‘little brain’. Development, 141(21), 4031–4041. 10.1242/dev.106559.

10. Chen, S., Zhong, X., Jiang, L., Zheng, X., Xiong, Y., Ma, S., Qiu, M., Huo, S., Ge, J., & Chen, Q. (2016). Maternal autoimmune diseases and the risk of autism spectrum disorders in offspring: A systematic review and meta-analysis. Behavioural Brain Research, 296, 61–69. 10.1016/j.bbr.2015.08.035.

11. Comi, A. M., Zimmerman, A. W., Frye, V. H., Law, P. A., & Peeden, J. N. (1999). Familial clustering of autoimmune disorders and evaluation of medical risk factors in autism. Journal of Child Neurology, 14(6), 388–394.

12. Courchesne, E., Pierce, K., Schumann, C. M., Redcay, E., Buckwalter, J. A., Kennedy, D. P., & Morgan, J. (2007). Mapping Early Brain Development in Autism. Neuron, 56(2), 399–413. 10.1016/j.neuron.2007.10.016.

13. Crawley, J. N. (2004). Designing mouse behavioral tasks relevant to autistic-like behaviors. Mental Retardation and Developmental Disabilities Research Reviews, 10(4), 248–258. 10.1002/mrdd.20039.

14. Crawley, J. N. (2007). What’s wrong with my mouse?: Behavioral phenotyping of transgenic and knockout mice (2nd ed.). Wiley-Interscience.

15. Dalakas, M. C. (2010). Pathogenesis and Treatment of Anti-MAG Neuropathy. Current Treatment Options in Neurology, 12(2), 71–83. 10.1007/s11940-010-0065-x.

16. Diamond, B., Huerta, P. T., Mina-Osorio, P., Kowal, C., & Volpe, B. T. (2009). Losing your nerves? Maybe it’s the antibodies. Nature Reviews Immunology, 9(6), 449–456. 10.1038/nri2529.

17. D’Mello, A. M., Crocetti, D., Mostofsky, S. H., & Stoodley, C. J. (2015). Cerebellar gray matter and lobular volumes correlate with core autism symptoms. NeuroImage: Clinical, 7, 631–639. 10.1016/j.nicl.2015.02.007.

18. D’Mello, A. M., & Stoodley, C. J. (2015). Cerebro-cerebellar circuits in autism spectrum disorder. Frontiers in Neuroscience, 9. 10.3389/fnins.2015.00408.

19. Domeniconi, M., Cao, Z., Spencer, T., Sivasankaran, R., Wang, K. C., Nikulina, E., Kimura, N., Cai, H., Deng, K., Gao, Y., He, Z., & Filbin, M. T. (2002). Myelin-Associated Glycoprotein Interacts with the Nogo66 Receptor to Inhibit Neurite Outgrowth. Neuron, 35(2), 283–290. 10.1016/S0896-6273(02)00770-5.

20. Estes, M. L., & McAllister, A. K. (2015). Immune mediators in the brain and peripheral tissues in autism spectrum disorder. Nature Reviews Neuroscience, 16(8), 469–486. (108429990). 10.1038/nrn3978.

21. Fatemi, S. H., Aldinger, K. A., Ashwood, P., Bauman, M. L., Blaha, C. D., Blatt, G. J., Chauhan, A., Chauhan, V., Dager, S. R., Dickson, P. E., Estes, A. M., Goldowitz, D., Heck, D. H., Kemper, T. L., King, B. H., Martin, L. A., Millen, K. J., Mittleman, G., Mosconi, M. W., … Welsh, J. P. (2012). Consensus Paper: Pathological Role of the Cerebellum in Autism. The Cerebellum, 11(3), 777–807. 10.1007/s12311-012-0355-9.

22. Fatemi, S. H., Halt, A. R., Realmuto, G., Earle, J., Kist, D. A., Thuras, P., & Merz, A. (2002). Purkinje Cell Size Is Reduced in Cerebellum of Patients with Autism. Cellular and Molecular Neurobiology, 22(2), 171–175. 10.1023/A:1019861721160.

23. Fernández-Suárez, D., Krapacher, F. A., Andersson, A., Ibáñez, C. F., & Kisiswa, L. (2019). MAG induces apoptosis in cerebellar granule neurons through p75NTR demarcating granule layer/white matter boundary. Cell Death & Disease, 10(10), 732. 10.1038/s41419-019-1970-x.

24. Fields, R. D. (2008). White matter in learning, cognition and psychiatric disorders. Trends in Neurosciences, 31(7), 361–370. 10.1016/j.tins.2008.04.001.

25. Fields, R. D., Woo, D. H., & Basser, P. J. (2015). Glial Regulation of the Neuronal Connectome through Local and Long-Distant Communication. Neuron, 86(2), 374–386. 10.1016/j.neuron.2015.01.014.

26. Gal, A., Raykin, E., Giladi, S., Lederman, D., Kofman, O., & Golan, H. M. (2023). Temporal dynamics of isolation calls emitted by pups in environmental and genetic mouse models of autism spectrum disorder. Frontiers in Neuroscience, 17. 10.3389/fnins.2023.1274039.

27. Geschwind, D. H. (2009). Advances in Autism. Annual Review of Medicine, 60(Volume 60, 2009), 367–380. 10.1146/annurev.med.60.053107.121225.

28. Geschwind, D. H., & Levitt, P. (2007). Autism spectrum disorders: Developmental disconnection syndromes. Current Opinion in Neurobiology, 17(1), 103–111. 10.1016/j.conb.2007.01.009

29. Govek, E.-E., Newey, S. E., & Aelst, L. V. (2005). The role of the Rho GTPases in neuronal development. Genes & Development, 19(1), 1–49. 10.1101/gad.1256405.

30. Hatten, M. E., & Heintz, N. (1995). Mechanisms of Neural Patterning and Specification in the Development Cerebellum. Annual Review of Neuroscience, 18(Volume 18, 1995), 385–408. 10.1146/annurev.ne.18.030195.002125.

31. Latov, N. (2014). Diagnosis and treatment of chronic acquired demyelinating polyneuropathies. Nature Reviews. Neurology, 10(8), 435–446. 10.1038/nrneurol.2014.117.

32. Li, C., Tropak, M. B., Gerlai, R., Clapoff, S., Abramow-Newerly, W., Trapp, B., Peterson, A., & Roder, J. (1994). Myelination in the absence of myelin-associated glycoprotein. Nature, 369(6483), 747–750. 10.1038/369747a0.

33. Lopez, P. H. H. (2014). Role of Myelin-Associated Glycoprotein (Siglec-4a) in the Nervous System. Advances in Neurobiology, 9, 245–262. 10.1007/978-1-4939-1154-7_11.

34. Lopez, P. H. H., & Báez, B. B. (2018). Gangliosides in Axon Stability and Regeneration. En Progress in Molecular Biology and Translational Science (Vol. 156, pp. 383–412). Elsevier. 10.1016/bs.pmbts.2018.03.001.

35. Lopez, P. H., Zhang, G., Zhang, J., Lehmann, H. C., Griffin, J. W., Schnaar, R. L., & Sheikh, K. A. (2010). Passive Transfer of IgG Anti-GM1 Antibodies Impairs Peripheral Nerve Repair. Journal of Neuroscience, 30(28), 9533–9541. 10.1523/JNEUROSCI.2281-10.2010.

36. Lord, C., Elsabbagh, M., Baird, G., & Veenstra-Vanderweele, J. (2018). Autism spectrum disorder. Lancet, 392(10146), 508–520. 10.1016/S0140-6736(18)31129-2.

37. Malkova, N. V., Yu, C. Z., Hsiao, E. Y., Moore, M. J., & Patterson, P. H. (2012). Maternal immune activation yields offspring displaying mouse versions of the three core symptoms of autism. Brain, Behavior, and Immunity, 26(4), 607–616. 10.1016/j.bbi.2012.01.011.

38. Martínez-Cerdeño, V., Camacho, J., Fox, E., Miller, E., Ariza, J., Kienzle, D., Plank, K., Noctor, S. C., & Van de Water, J. (2016). Prenatal Exposure to Autism-Specific Maternal Autoantibodies Alters Proliferation of Cortical Neural Precursor Cells, Enlarges Brain, and Increases Neuronal Size in Adult Animals. Cerebral Cortex, 26(1), 374–383. 10.1093/cercor/bhu291.

39. Meyer-Franke, A., Tropak, M. B., Roder, J. C., Fischer, P., Beyreuther, K., Probstmeier, R., & Schachner, M. (1995). Functional topography of myelin-associated glycoprotein. II. Mapping of domains on molecular fragments. Journal of Neuroscience Research, 41(3), 311–323. 10.1002/jnr.490410304.

40. Mi, S., Lee, X., Shao, Z., Thill, G., Ji, B., Relton, J., Levesque, M., Allaire, N., Perrin, S., Sands, B., Crowell, T., Cate, R. L., McCoy, J. M., & Pepinsky, R. B. (2004). LINGO-1 is a component of the Nogo-66 receptor/p75 signaling complex. Nature Neuroscience, 7(3), 221–228. (12345232). 10.1038/nn1188.

41. Mostafa, G. A., & Al-Ayadhi, L. Y. (2013). The possible relationship between allergic manifestations and elevated serum levels of brain specific auto-antibodies in autistic children. Journal of Neuroimmunology, 261(1), 77–81. 10.1016/j.jneuroim.2013.04.003.

42. Mostafa, G. A., Awad El-Sayed, Z., Mohamed Abd El-Aziz, M., & Farouk El-Sayed, M. (2008). Serum Anti-Myelin—Associated Glycoprotein Antibodies in Egyptian Autistic Children. Journal of Child Neurology, 23(12), 1413–1418. 10.1177/0883073808319321.

43. Mount, C. W., & Monje, M. (2017). Wrapped to Adapt: Experience-Dependent Myelination. Neuron, 95(4), 743–756. 10.1016/j.neuron.2017.07.009

44. Moy, S. S., Nadler, J. J., Perez, A., Barbaro, R. P., Johns, J. M., Magnuson, T. R., Piven, J., & Crawley, J. N. (2004). Sociability and preference for social novelty in five inbred strains: An approach to assess autistic-like behavior in mice. *Genes*, Brain and Behavior, 3(5), 287–302. 10.1111/j.1601-1848.2004.00076.x.

45. Mulherkar, S., Uddin, M. D., Couvillon, A. D., Sillitoe, R. V., & Tolias, K. F. (2014). The small GTPases RhoA and Rac1 regulate cerebellar development by controlling cell morphogenesis, migration and foliation. Developmental Biology, 394(1), 39–53. 10.1016/j.ydbio.2014.08.004.

46. Nguyen, T., Mehta, N. R., Conant, K., Kim, K.-J., Jones, M., Calabresi, P. A., Melli, G., Hoke, A., Schnaar, R. L., Ming, G.-L., Song, H., Keswani, S. C., & Griffin, J. W. (2009). Axonal protective effects of the myelin-associated glycoprotein. The Journal of Neuroscience: The Official Journal of the Society for Neuroscience, 29(3), 630–637. 10.1523/JNEUROSCI.5204-08.2009.

47. Palandri, A., Salvador, V. R., Wojnacki, J., Vivinetto, A. L., Schnaar, R. L., & Lopez, P. H. H. (2015). Myelin-associated glycoprotein modulates apoptosis of motoneurons during early postnatal development via NgR/p75NTR receptor-mediated activation of RhoA signaling pathways. Cell Death and Disease, 6, e1876. 10.1038/cddis.2015.228.

48. Palay, S. L. (with Chan-Palay, V.). (1974). Cerebellar cortex: Cytology and organization. Springer.

49. Pan, B., Fromholt, S. E., Hess, E. J., Crawford, T. O., Griffin, J. W., Sheikh, K. A., & Schnaar, R. L. (2005). Myelin-associated glycoprotein and complementary axonal ligands, gangliosides, mediate axon stability in the CNS and PNS: Neuropathology and behavioral deficits in single- and double-null mice. Experimental Neurology, 195(1), 208–217. 10.1016/j.expneurol.2005.04.017.

50. Patterson, P. H. (2011). Maternal infection and immune involvement in autism. Trends in Molecular Medicine, 17(7), 389–394. 10.1016/j.molmed.2011.03.001.

51. Peça, J., Feliciano, C., Ting, J. T., Wang, W., Wells, M. F., Venkatraman, T. N., Lascola, C. D., Fu, Z., & Feng, G. (2011). Shank3 mutant mice display autistic-like behaviours and striatal dysfunction. Nature, 472(7344), 437–442. 10.1038/nature09965.

52. Pernet, V., Joly, S., Christ, F., Dimou, L., & Schwab, M. E. (2008). Nogo-A and Myelin-Associated Glycoprotein Differently Regulate Oligodendrocyte Maturation and Myelin Formation. Journal of Neuroscience, 28(29), 7435–7444. 10.1523/JNEUROSCI.0727-08.2008.

53. Poltorak, M., Sadoul, R., Keilhauer, G., Landa, C., Fahrig, T., & Schachner, M. (1987). Myelin-Associated Glycoprotein, a Member of the L2/HNK-1 Family of Neural Cell Adhesion Molecules, Is Involved in Neuron-Oligodendrocyte and Oligodendrocyte-Oligodendrocyte Interaction. The Journal of Cell Biology, 105(4), 1893–1899.

54. Rozas, G., Guerra, M. J., & Labandeira-García, J. L. (1997). An automated rotarod method for quantitative drug-free evaluation of overall motor deficits in rat models of parkinsonism. Brain Research Protocols, 2(1), 75–84. 10.1016/S1385-299X(97)00034-2.

55. Sakurai, T., Ramoz, N., Barreto, M., Gazdoiu, M., Takahashi, N., Gertner, M., Dorr, N., Gama Sosa, M. A., De Gasperi, R., Perez, G., Schmeidler, J., Mitropoulou, V., Le, H. C., Lupu, M., Hof, P. R., Elder, G. A., & Buxbaum, J. D. (2010). *Slc25a12* Disruption Alters Myelination and Neurofilaments: A Model for a Hypomyelination Syndrome and Childhood Neurodevelopmental Disorders. Biological Psychiatry, Metabotropic Glutamate Receptor Agonist Modulation of Alcohol Consumption, 67(9), 887–894. 10.1016/j.biopsych.2009.08.042.

56. Sandin, S., Lichtenstein, P., Kuja-Halkola, R., Hultman, C., Larsson, H., & Reichenberg, A. (2017). The Heritability of Autism Spectrum Disorder. JAMA, 318(12), 1182–1184. 10.1001/jama.2017.12141.

57. Scattoni, M. L., Gandhy, S. U., Ricceri, L., & Crawley, J. N. (2008). Unusual Repertoire of Vocalizations in the BTBR T+tf/J Mouse Model of Autism. PLOS ONE, 3(8), e3067. 10.1371/journal.pone.0003067.

58. Schmued, L. C., Stowers, C. C., Scallet, A. C., & Xu, L. (2005). Fluoro-Jade C results in ultra high resolution and contrast labeling of degenerating neurons. Brain Research, 1035(1), 24–31. 10.1016/j.brainres.2004.11.054.

59. Semenova, M. M., Mäki-Hokkonen, A. M. J., Cao, J., Komarovski, V., Forsberg, K. M., Koistinaho, M., Coffey, E. T., & Courtney, M. J. (2007). Rho mediates calcium-dependent activation of p38α and subsequent excitotoxic cell death. Nature Neuroscience, 10(4), 436–443. (24486829). 10.1038/nn1869.

60. Shair, H. N. (2007). Acquisition and expression of a socially mediated separation response. *Behavioural Brain Research, Mammalian Vocalization: Neural*, Behavioural, and Environmental Determinants, 182(2), 180–192. 10.1016/j.bbr.2007.02.016.

61. Sillitoe, R. V., & Joyner, A. L. (2007). Morphology, Molecular Codes, and Circuitry Produce the Three-Dimensional Complexity of the Cerebellum. Annual Review of Cell and Developmental Biology, 23(Volume 23, 2007), 549–577. 10.1146/annurev.cellbio.23.090506.123237.

62. Silverman, J. L., Yang, M., Lord, C., & Crawley, J. N. (2010). Behavioural phenotyping assays for mouse models of autism. Nature reviews. Neuroscience, 11(7), 490–502. 10.1038/nrn2851.

63. Singer, H. S., Morris, C. M., Williams, P. N., Yoon, D. Y., Hong, J. J., & Zimmerman, A. W. (2006). Antibrain antibodies in children with autism and their unaffected siblings. Journal of Neuroimmunology, 178(1-2), 149–155. 10.1016/j.jneuroim.2006.05.025.

64. Skefos, J., Cummings, C., Enzer, K., Holiday, J., Weed, K., Levy, E., Yuce, T., Kemper, T., & Bauman, M. (2014). Regional Alterations in Purkinje Cell Density in Patients with Autism. PLOS ONE, 9(2), e81255. 10.1371/journal.pone.0081255.

65. Sotelo, C. (2004). Cellular and genetic regulation of the development of the cerebellar system. Progress in Neurobiology, 72(5), 295–339. 10.1016/j.pneurobio.2004.03.004.

66. State, M. W., & Šestan, N. (2012). Neuroscience. The emerging biology of autism spectrum disorders. Science, 337(6100), 1301–1303. 10.1126/science.1224989.

67. Steck, A. J. (2021). Anti-MAG neuropathy: From biology to clinical management. Journal of Neuroimmunology, 361, 577725. 10.1016/j.jneuroim.2021.577725.

68. Tsai, P. T., Hull, C., Chu, Y., Greene-Colozzi, E., Sadowski, A. R., Leech, J. M., Steinberg, J., Crawley, J. N., Regehr, W. G., & Sahin, M. (2012). Autistic-like behaviour and cerebellar dysfunction in Purkinje cell Tsc1 mutant mice. Nature, 488(7413), 647–651. (79448502). 10.1038/nature11310.

69. Vojdani, A., Campbell, A. W., Anyanwu, E., Kashanian, A., Bock, K., & Vojdani, E. (2002). Antibodies to neuron-specific antigens in children with autism: Possible cross-reaction with encephalitogenic proteins from milk, *Chlamydia pneumoniae* and *Streptococcus* group A. Journal of Neuroimmunology, 129(1), 168–177. 10.1016/S0165-5728(02)00180-7.

70. Wang, S. S.-H., Kloth, A. D., & Badura, A. (2014). The Cerebellum, Sensitive Periods, and Autism. Neuron, 83(3), 518–532. 10.1016/j.neuron.2014.07.016.

71. Wang, V. Y., & Zoghbi, H. Y. (2001). Genetic regulation of cerebellar development. Nature Reviews Neuroscience, 2(7), 484–491. (11360035). 10.1038/35081558.

72. Wang, Y., Cao, A., Wang, J., Bai, H., Liu, T., Sun, C., Li, Z., Tang, Y., Xu, F., & Liu, S. (2025). Abnormalities in cerebellar subregions’ volume and cerebellocerebral structural covariance in autism spectrum disorder. Autism Research, 18(1), 83–97. 10.1002/aur.3287.

73. White, J. J., & Sillitoe, R. V. (2013). Postnatal development of cerebellar zones revealed by neurofilament heavy chain protein expression. Frontiers in Neuroanatomy, 7, 9. 10.3389/fnana.2013.00009.

74. Wills, S., Cabanlit, M., Bennett, J., Ashwood, P., Amaral, D., & Van de Water, J. (2007). Autoantibodies in autism spectrum disorders (ASD). Annals of the New York Academy of Sciences, 1107, 79–91. 10.1196/annals.1381.009.

75. Yakovlev PI. (1967). The myelogenetic cycles of regional maturation of the brain. Regional development of the brain in early life, 3–70.

76. Yin, X., Chen, L., Xia, Y., Cheng, Q., Yuan, J., Yang, Y., Wang, Z., Wang, H., Dong, J., Ding, Y., & Zhao, X. (2016). Maternal Deprivation Influences Pup Ultrasonic Vocalizations of C57BL/6J Mice. PLOS ONE, 11(8), e0160409. 10.1371/journal.pone.0160409.

77. Yiu, G., He, Z., & He, Z. (2006). Glial inhibition of CNS axon regeneration. Nature Reviews Neuroscience, 7(8), 617–627. (22155263). 10.1038/nrn1956.

