## Supplementary figures and images for "Disruption of Myelin-Associated Glycoprotein Activity Drives Aberrant Cerebellar Neurodevelopment and Autism-Like Behaviors"

### Suppl Figure 1

#### S1- Supplementary Figure 1

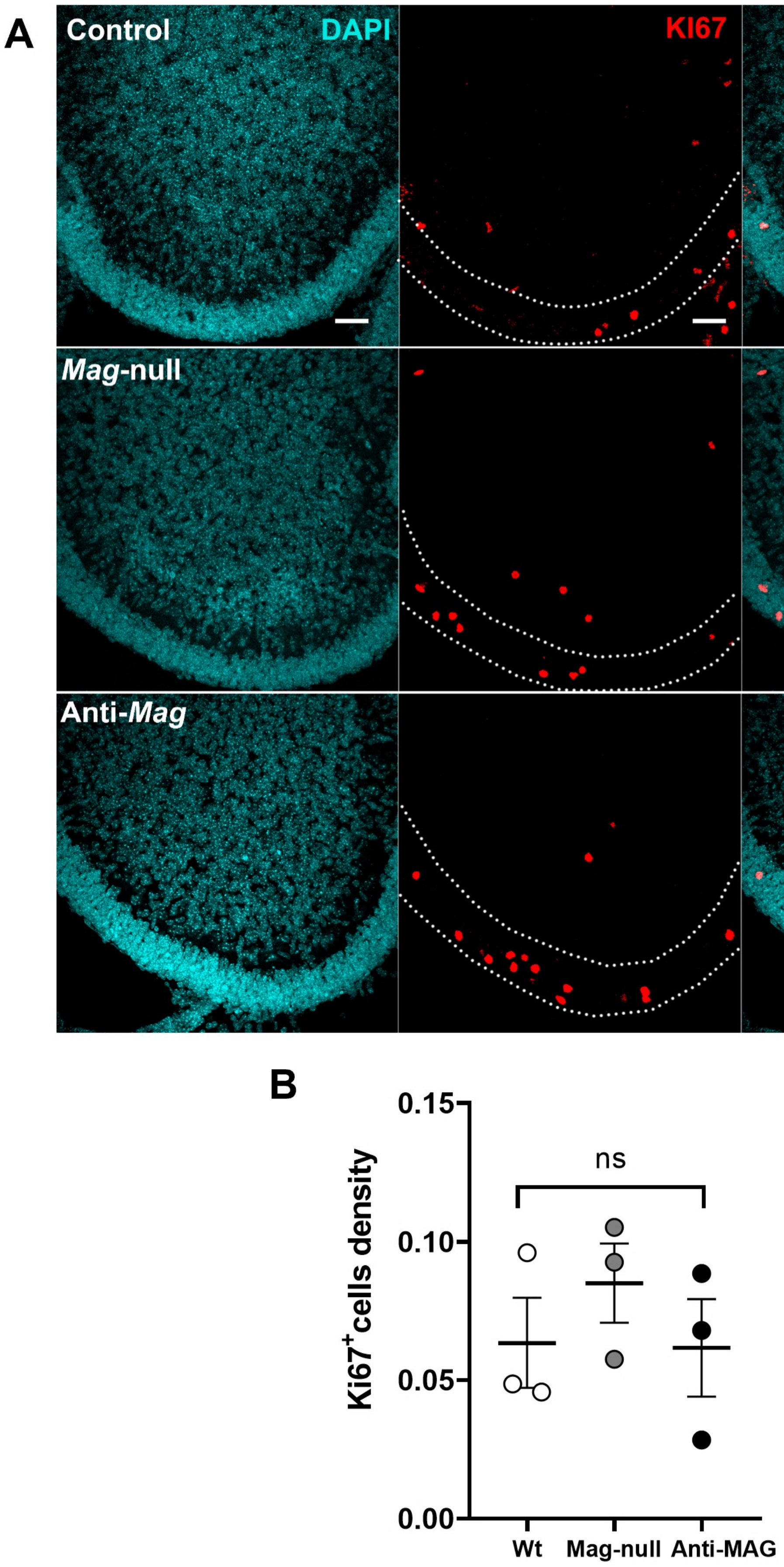

### Suppl Figure 2

S2-Supplementary Figure 2

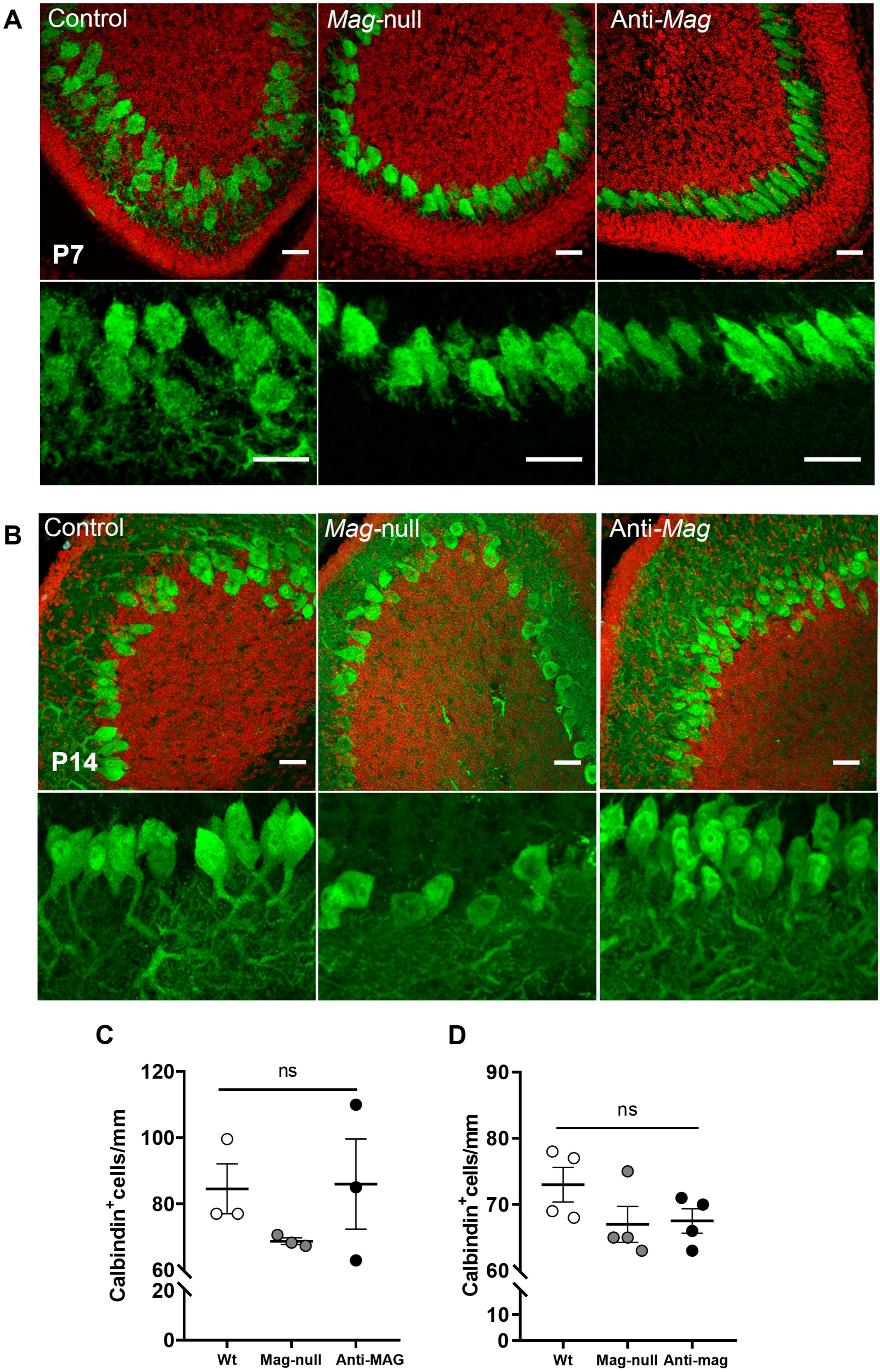
